# Quantifying Distributions of Cryo-EM Projections

**DOI:** 10.1101/2025.08.14.670280

**Authors:** Alexandre G. Urzhumtsev

## Abstract

In cryo-electron microscopy, a set of two-dimensional projections collected from different viewing directions may complicate image processing and subsequent model building if the distribution of these views is non-uniform. View distributions are traditionally represented as color-coded two-dimensional diagrams. However, such diagrams can introduce distortions, are cumbersome to manipulate, store, and compare across datasets; they do not provide a commonly accepted quantitative measure of uniformity. In this work, we propose a method to characterize these angular distributions quantitatively and to represent them as simple one-dimensional curves rather than two-dimensional colored diagrams. The suggested measures could be incorporated into databases such as the EMDB to provide a compact, standardized description of the angular distribution of particle views.

**Synopsis:** We propose a numerical measure to quantify the deviation of a cryo-EM 2D view distribution from the uniform one. This approach allows the traditional two-dimensional projection diagrams to be replaced, or completed with simple, easily generated curves that are straightforward to analyze.

## 1. Introduction

In cryo-electron microscopy (cryo-EM), three-dimensional (3D) reconstructions are derived from two-dimensional (2D) projections of the specimen. The spatial distribution of these projections is one of critical factors in determining the success of the reconstruction. Strongly overrepresented, underrepresented, or entirely missing views can render the resulting 3D map difficult to interpret (*e.g*., Unger, 2000; Sorzano *et al*., 2021; and many others).

Each orthogonal projection is characterized by its direction, or equivalently by the intersection of the projection axis with the surface of a unit sphere centered on the object. The distribution of these directions, referred to as *views*, can be visualized as a relief map on the spherical surface, as implemented, for example, through the combined use of *Relion* (Scheres, 2012) and *Chimera* (Pettersen et al., 2004). For practical purposes and in publications, this spherical distribution is typically displayed as a two-dimensional projection, often color-coded to indicate views density (*e.g*., Orlov *et al*., 2006; Punjani *et al*., 2017; Grant *et al*., 2018). However, most such projection methods do not preserve the surface area of spherical elements, potentially distorting the perceived distribution. To fix this problem, we recently developed the program *VUE* (Urzhumtseva *et al*., 2024), which employs the Lambert projection.

While such 2D representations, plotted with respect to a chosen reference direction, provide a visual and qualitative impression of the views distribution, they are not straightforward to characterize these distributions numerically, to be compared, reported, or archived. Moreover, they do not yield a single quantitative measure indicating how far a given distribution deviates from the uniform one.

The aim of the present work is to simplify these 2D representations by reducing them to one-dimensional curves, thereby making the information more easily comparable and transferable. Furthermore, we propose single numerical descriptors of distribution non-uniformity, which can serve as a concise reference value for reporting and database annotation.

In the following, 2D views distributions are displayed using the *“individual views”* option of the *VUE* program. Views frequencies are color codes, with the values expressed in a logarithmic scheme defined as log_2_(*view_frequency* / *mean_frequency*). For direct comparability, the same color scale is applied consistently across all 2D diagrams.

The experimental data sets used in this study stem from recent experimental cryo-EM data (acquired on the onsite Titan Krios G4 cryo electron microscope installed at the CBI) obtained in the laboratory when working on various ribosomal structures and complexes.

## 2. Method

### 2.1 Theoretical uniform distribution

Let’s choose the origin of the coordinate system, point **O**, at the center of the object of interest. The direction of each orthogonal projection, *i.e*., a view of such object, can be represented by the intersection of its projection line with the unit sphere centered at **O**. The distribution of views therefore corresponds to the distribution of points on the surface of this sphere. While the problem of distributing points uniformly on a spherical surface is a classical one, extensively discussed in the literature and online, the characterization of such distributions has received comparatively less attention (see, for example, Del Bono *et al.*, 2024, and references therein).

The essential point here is that, when analyzing a distribution of points on a sphere, we are concerned with the distances between points rather than their absolute positions with respect to a chosen direction, that orthogonal to the page. This perspective is consistent with the considerations of Del Bono *et al.* (2024). However, instead of investigating a formal potential between points, we propose a simpler purely geometric approach.

First, since two opposite views in cryo-EM contain identical information, each point in the southern hemisphere is replaced, if present, by its antipodal point in the northern hemisphere. Let *d* denote the geodesic distance between two points on the unit sphere, measured along the great-circle arc connecting them. This distance is numerically equal to the central angle *δ*, expressed in radians. In particular, for a reference point at the North Pole (**N**), the angular distance *δ* to any other point **M** coincides with the zenith angle *θ*, that is, the angle between the vectors **ON** and **OM** (see, *e.g*., Urzhumtsev & Urzhumtseva, 2019, for definitions). For points in the northern hemisphere, 0 ≤ θ ≤ *π*/2.

If the sphere is uniformly covered by a sufficiently large number of points, the number of points lying on the circle defined by a given angle θ is proportional to sin θ. Interpreting these points as randomly distributed, the normalized probability density for the distance from **N** to all points on the hemisphere is

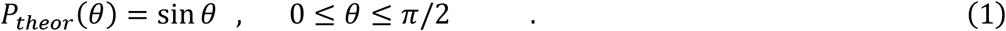

Since all points on the sphere are equivalent under a uniform distribution, the point-to-point distance distribution is identical for any chosen reference point. Consequently, integrating over all uniformly distributed views yields the same expression (1) for the total probability density of the point-to-point distance. The corresponding cumulative distribution function is

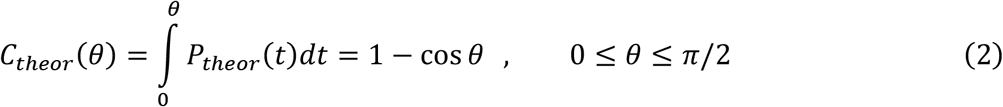

### 2.2 Numerical simulation of the uniform distribution

For a discrete set of views (points **M**_k_ on the sphere), we proceed as follows. First, all pairwise geodesic distances are computed. A histogram of these distances is then constructed and normalized to yield the corresponding relative frequencies, *f*_*n*_, such that

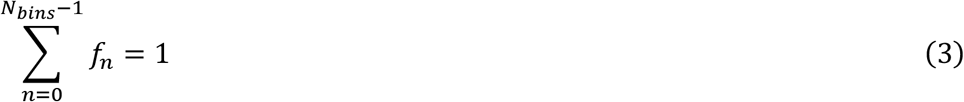

Here the bin boundaries are chosen uniformly,

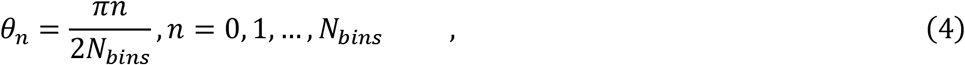

and the distance between **M**_j_ and **M**_k_ is calculated as the arccosine of the dot product of the corresponding unit vectors **OM**_j_ and **OM**_k_. The resulting histogram is then converted into a probability distribution by normalizing the frequencies

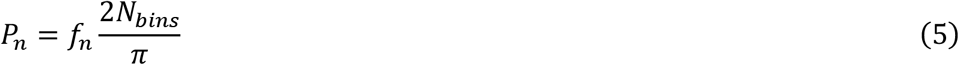

Yielding

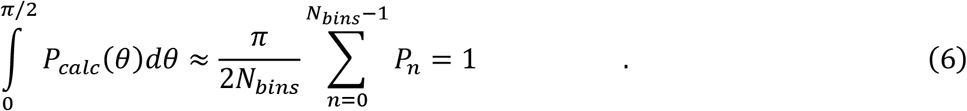

A wide range of statistical tools exists for comparing numerically calculated frequencies *f*_*n*_ with their theoretical counterparts. In our case, however, the number of projections is both very large and varies substantially between structures, while our main interest lies in comparing results across different structures. For this reason, we do not employ standard measures such as χ_2_or the Kolmogorov–Smirnov statistic (e.g., https://en.wikipedia.org/wiki/Kolmogorov-Smirnov_test), which explicitly depend on the number of measurements. Instead, we compute three numerical characteristics based on the core principles of these methods. The first of these is the root-mean-squared deviation (*rmsd*)

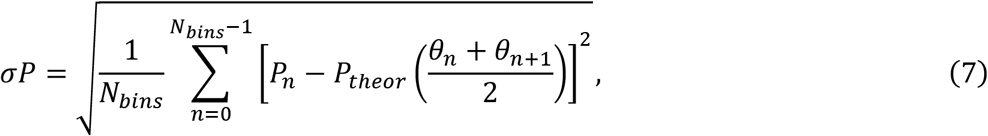

between the calculated and theoretical probability distributions. The second characteristic is the maximum absolute difference, across all bins, between the calculated and theoretical cumulative distributions.

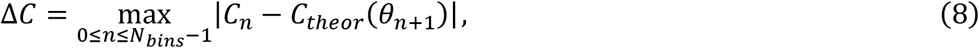

where

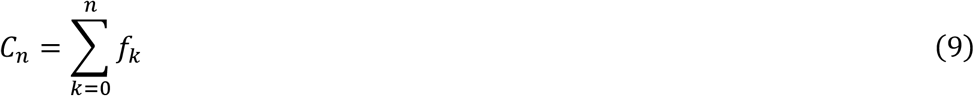

Both of these measures are integrative and therefore less sensitive to the choice of bin boundaries than local measures, such as the third characteristic that we also compute

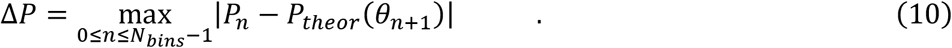

To compare Δ*P* values across different structures, it is necessary to use a common number of bins, *N*_bins_.

### 2.3. Parameters of the procedure

The computational cost of evaluating all point-to-point distances grows proportionally to the square of the number of views. To reduce CPU time, it is preferable to analyze a representative subset of views rather than the full set, provided that this does not significantly affect the results. This selection is intended solely for statistical analysis, not for subsequent 3D reconstruction, for which dedicated strategies have been discussed elsewhere (*e.g*., Efron *et al*., 2004). The subset is obtained by accepting or rejecting each projection in the original set with a given probability, uniformly distributed over the unit interval. The resulting subset size or, equivalently, the selection probability threshold, is a parameter that may influence the outcome.

Another factor affecting the results is the number of histogram bins. Too few bins give an overly coarse approximation to the continuous distribution, while too many bins may introduce artifacts in the frequency analysis due to insufficient counts per bin, particularly for small data sets.

To estimate suitable parameter values, we used a set of *N*_total_ = 100,000 views from our previous tests (Urzhumtseva *et al*., 2024) with simulated cryo-EM data for the IF2 atomic model (Simonetti *et al*., 2013). These views were generated randomly and uniformly, producing a uniform distribution of points on the unit sphere.

First, for all *N*_total_ views, we varied the number of intervals *N*_bins_, from several ten to several hundred, and compared the point-to-point distance distribution with the theoretical values as defined above by (7), (8), (10) as functions of *N*_bins_. The minimal error values, Δ*P* = 17 · 10^−5^ and σ*P* = 6 · 10^−5^, were obtained with *N*_bins_ = 60. These errors increased gradually with *N*_bins_, reaching Δ*P* = 50 · 10^−5^, σ*P* = 15 · 10^−5^ for *N*_bins_ = 360, and rose by roughly an order of magnitude for *N*_bins_ = 10. The difference between the cumulative functions Δ*C* remained essentially unchanged at about 10^−5^. For convenience to have one bin per angular degree, we used *N*_bins_ = 90 in all subsequent calculations. This choice, corresponding to a one-degree precision in point-to-point distances, also reflects the finite angular accuracy of projection annotations in practical studies.

Next, keeping the number of bins fixed at *N*_bins_ = 90, we progressively reduced the number of views, selecting the corresponding points to remain uniformly distributed on the spherical surface for each subset (**Fig. 1**). For these simulated, error-free sets of uniformly distributed views both Δ*P* and σ*P* were approximately inversely proportional to *N*_views_:

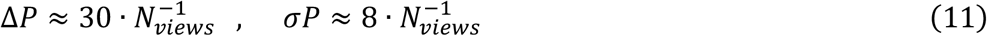

(details not shown). We assumed that working with approximately 5,000 - 10,000 views, allowing for reasonably fast calculations with basic laptops, the resulting errors would remain well below those arising from the intrinsic non-uniformity of angular view distributions. This assumption is to be validated using experimental data sets (see below).

**Figure 1.**
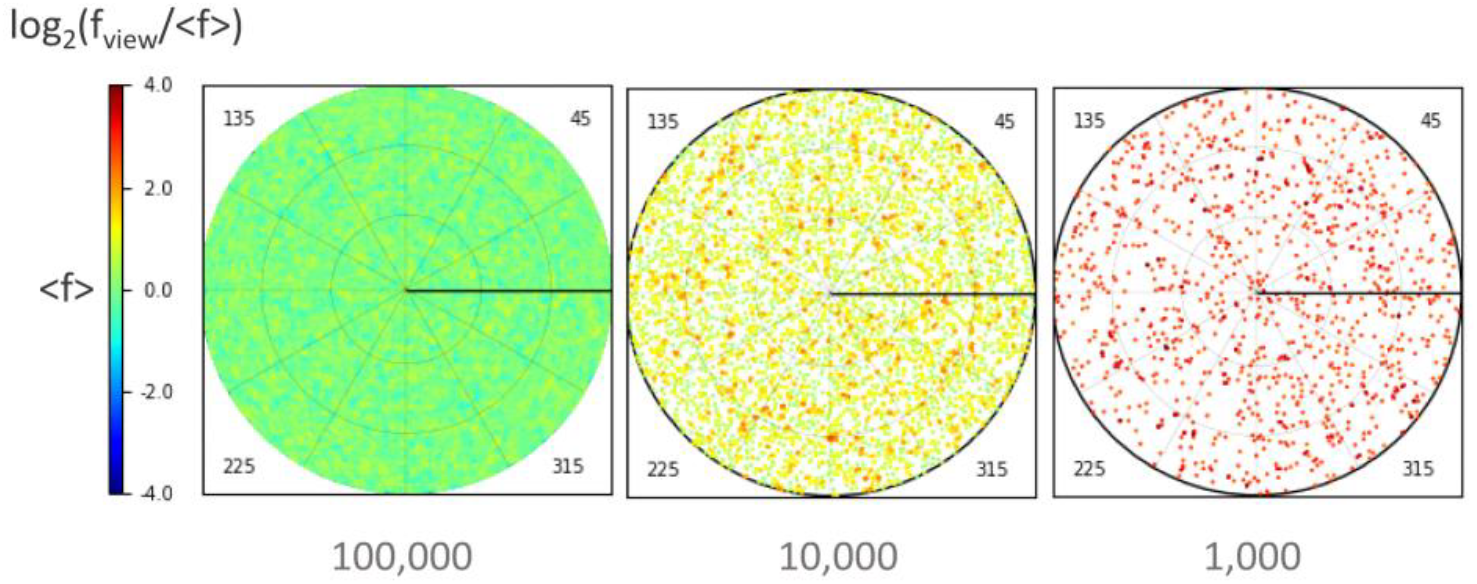
The 2D-diagrams (by program *VUE*) showing the Lambert projection of the points on the sphere, corresponding to the projection directions. Simulated views are uniformly distributed; the number of views is given under the diagrams. Frequencies are color-coded according to the logarithm of their ratio to the mean frequency; the same scheme is used for all other 2D-diagrams of the article.

### 2.4. Training with non-uniform simulated distributions

To generate reference point-to-point distance histograms and corresponding quantitative measures for non-uniform views distributions, we created several sets of points on the unit sphere. These points were normally distributed and clustered around one, two, or three centers positioned differently on the sphere. The parameters of the Gaussian distributions varied across sets. Each group consisted of 5,000 points. **Table 1** summarizes the parameters of these sets. **Fig. 2** presents the sets as 2D views diagrams, with the right column displaying the corresponding frequency curves *P*_*calc*_(θ) that represent the point-to-point distance distributions calculated as described above.

**Table 1.**
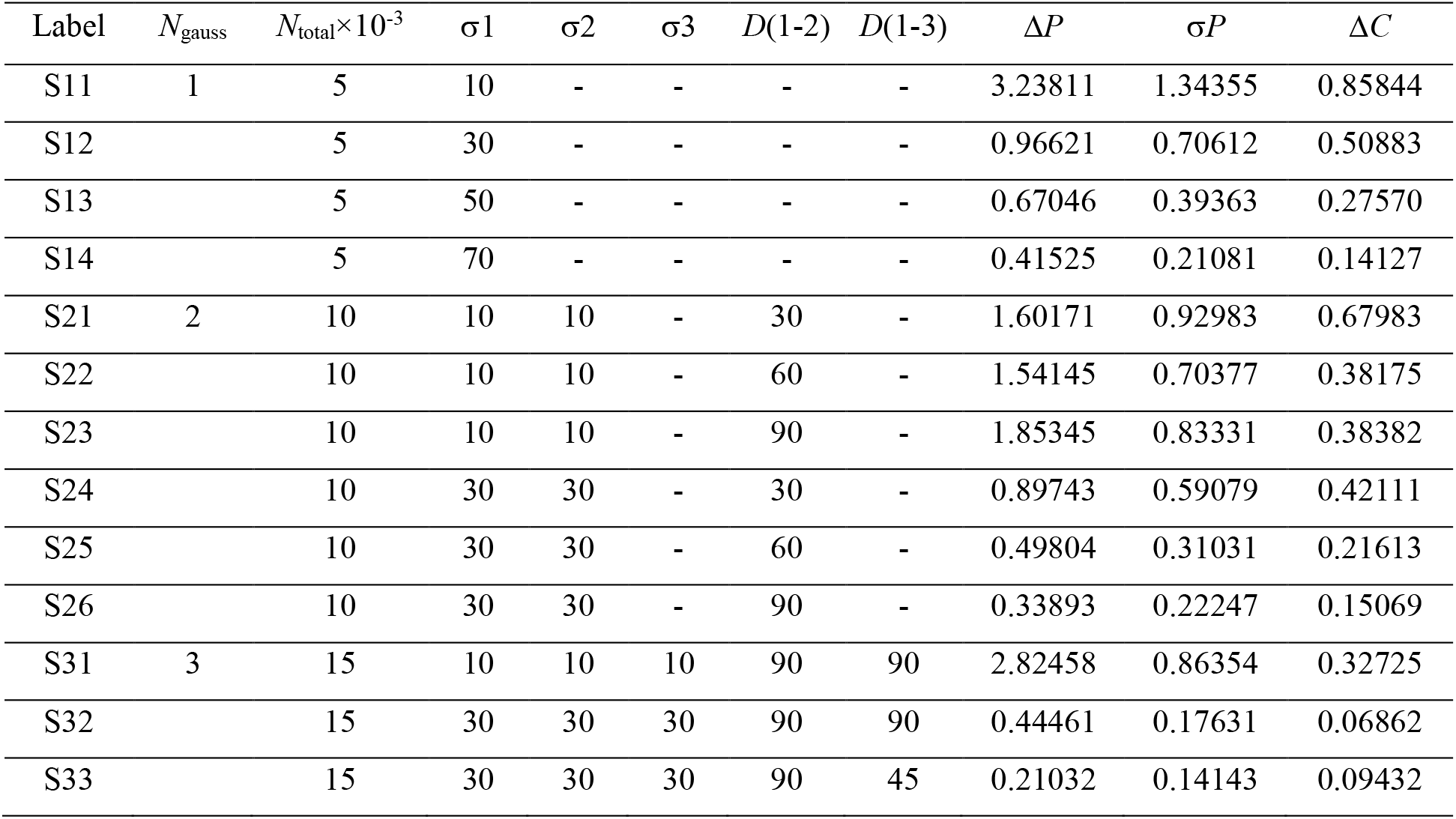
Point-to-point distance analysis for the simulated sets of views normally distributed around one, two, or three centers on the unit sphere. Parameters σn indicate the *rmsd* values for the Gaussian of the cluster *n*, each containing 5,000 views. *D*(*j*-*k*) shows the distance, in degrees, between the Gaussian centers *j* and *k*. Δ*P* (10), σ*P* (7), and ΔC (8) are the measures of the difference between the calculated and theoretical distributions as defined in the text.

**Figure 2.**
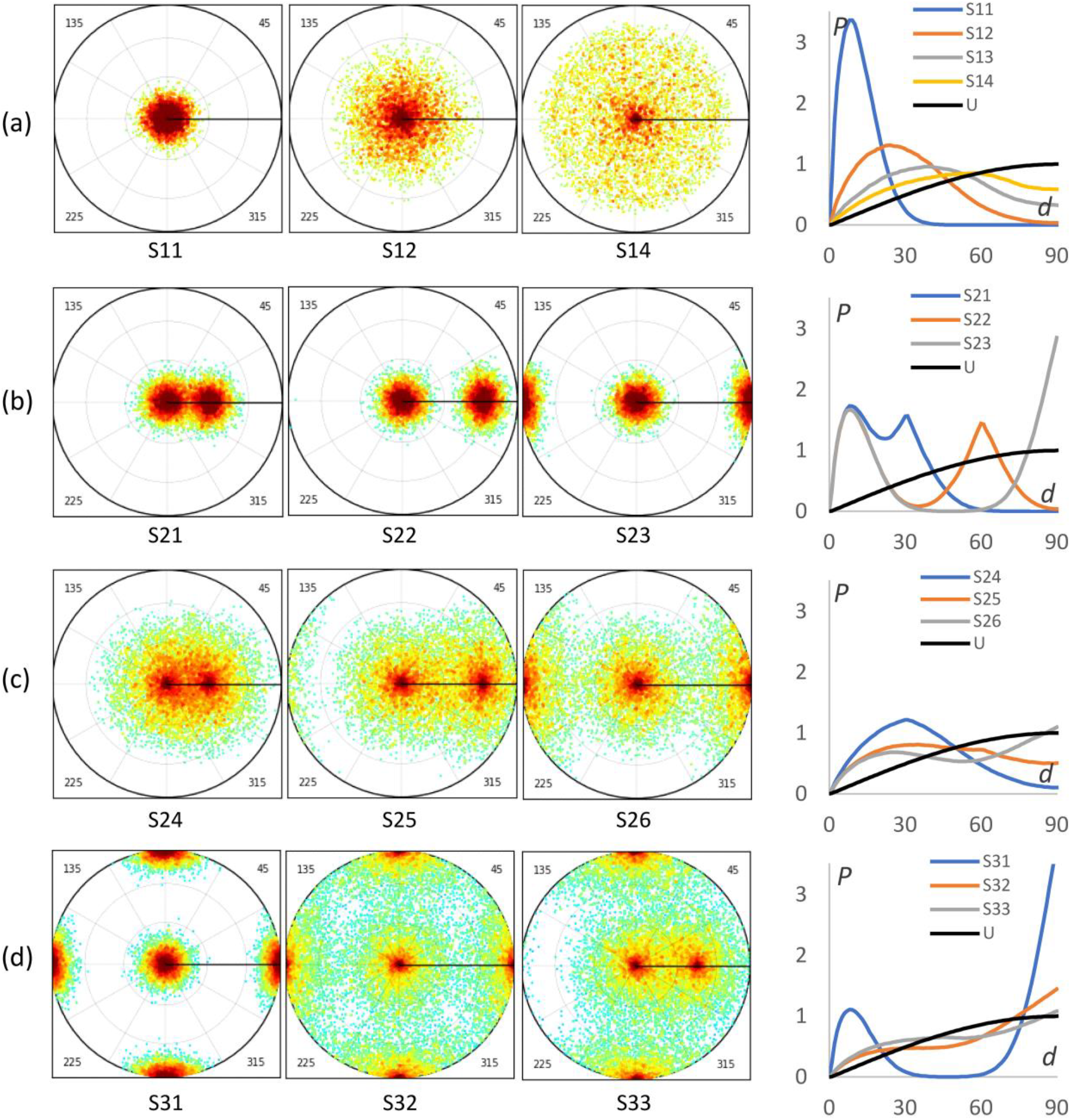
Simulated data sets normally distributed around several centers. Right column shows respective frequency curves. U (black curve) is theoretical for the uniform views distribution. Sets S*jk* contains the views randomly distributed around one (a), two (b, c) and three (d) centers with a different width and a different distance between centers (numerical values are given in Table 1).

The peaks in the frequency curves correspond to the clusters, reflecting their width and the distances between them, thereby condensing key information from the 2D distributions into a simple one-dimensional curve. Naturally, less detailed features are lost in this compression. As expected, the error values, Δ*P* and σ*P*, are several orders of magnitude larger than those calculated previously with the uniformly distributed data sets when reducing the size of the projection sets.

### 2.5. Cumulative function

When working with histograms and directly comparing the content of individual bins, measures like Δ*P* may introduce some artifacts. In contrast, integrative measures like σ*P* tend to be more robust. Another useful integrative measure, mentioned earlier, is the maximal difference between the cumulative distributions, ΔC, used in the Kolmogorov-Smirnov test. Initial calculations on simulated uniformly distributed views yielded ΔC values around 10^−4^, which remained largely independent of the number of bins, provided the bins were not too few, and stable as long as the number of views exceeded approximately 5,000. **Fig. 3** presents the cumulative distribution plots for the data sets shown in **Fig. 2**.

**Figure 3.**
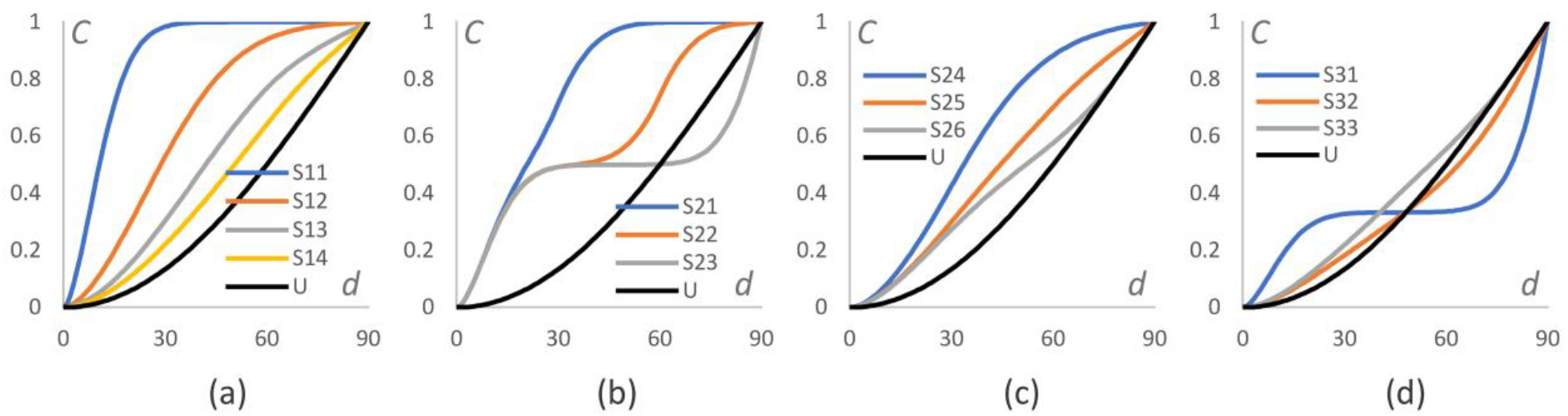
Cumulative function for the simulated data sets normally distributed around one, two, or three centers (see Fig. 2). Black curve (U) stands for the theoretical function for the uniform views distribution.

In particular, **Fig. 3a** demonstrates how the distribution of views approaches uniform one as the cluster width increases. Similarly, **Fig. 3c** shows this effect when the distance between cluster centers is increased. **Figs. 3b** and **3d** illustrate analogous trends for other sets of views.

The integrative measure ΔC is straightforwardly visualized using the cumulative distribution curves, where it corresponds to the maximal distance between them, much like Δ*P* relates to the frequency curves. In contrast, a simple geometric illustration of the integrative measure σ*P* is less intuitive.

## 3. Results: application to experimental data sets

### 3.1. Validation with experimental data

Before analyzing severely non-uniformly distributed experimental data sets, we aimed to validate our previous findings obtained from simulated data and to assess the robustness of the method. A key question was how effectively the frequency curves and associated statistical measures convey information about the spatial distribution of views without relying on 2D views diagrams.

To address this, we began by analyzing three independent experimental data sets corresponding to different samples of the same structure, hereafter referred to as ‘structure B’. Each set contained a relatively small number of projections, between 30,000 and 40,000 (**Table 2**), which allowed us to analyze the full sets of views.

**Table 2.**
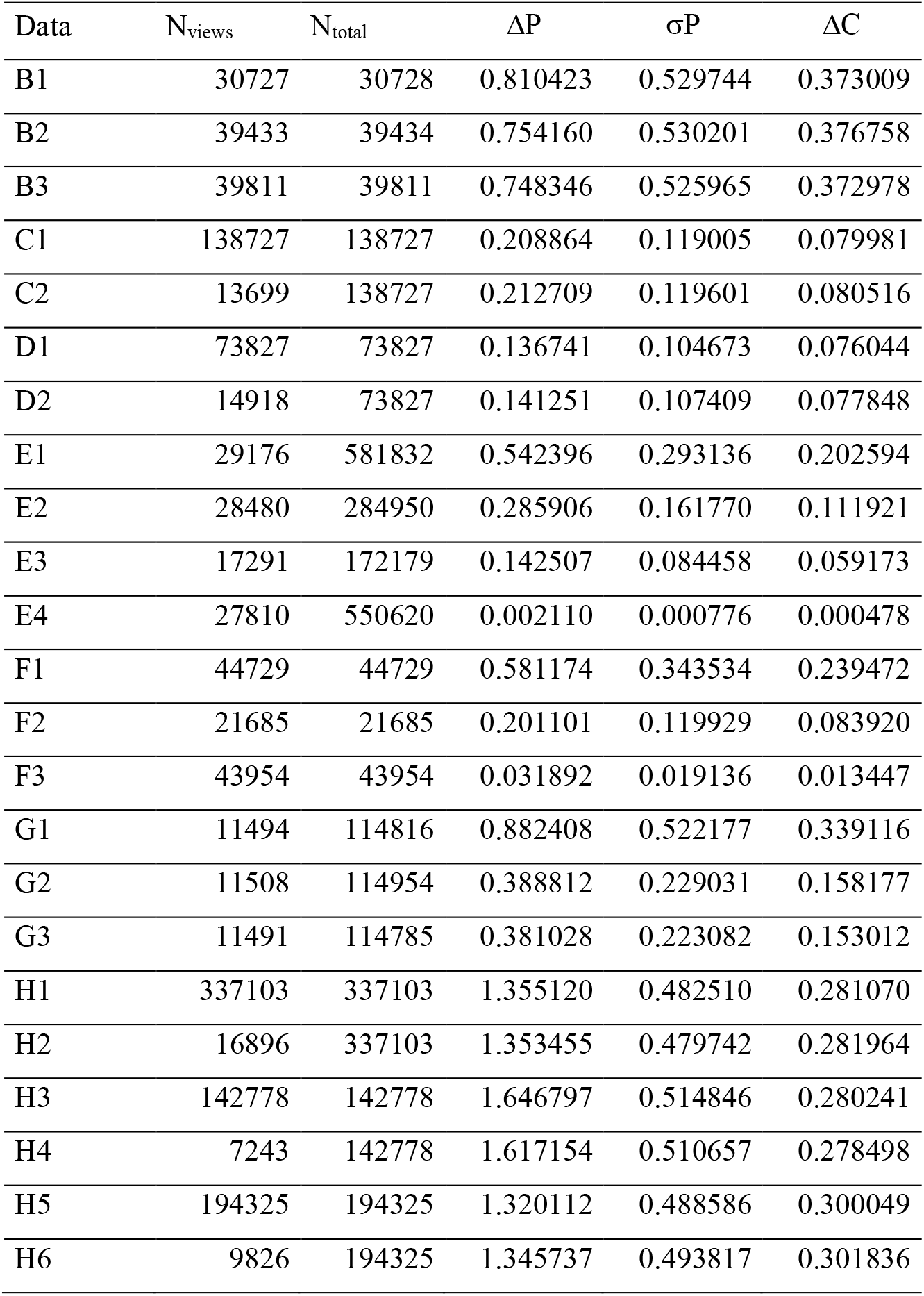
Point-to-point distance analysis for the experimental data sets. Δ*P* (10), σ*P* (7), and ΔC (8), defined in the text, express the difference between the calculated and uniform distributions.

The frequency curves for the three data sets were similar, each showing a single peak at approximately 30 degrees (**Fig. 2a**). This value aligns reasonably well with the Δ*P* values, which correspond to a Gaussian peak of about 40 degrees in width (compare with Δ*P* values for sets S12 and S13 in **Table 1**). The 2D diagrams of the views distributions confirm this (**Fig. 4a**). They show these views distributions being similar to each other by their shape, while with a slight difference in the peak position; this had no effect on the frequency curves nor on the overall quality of the angular distribution and subsequent reconstruction.

**Figure 4.**
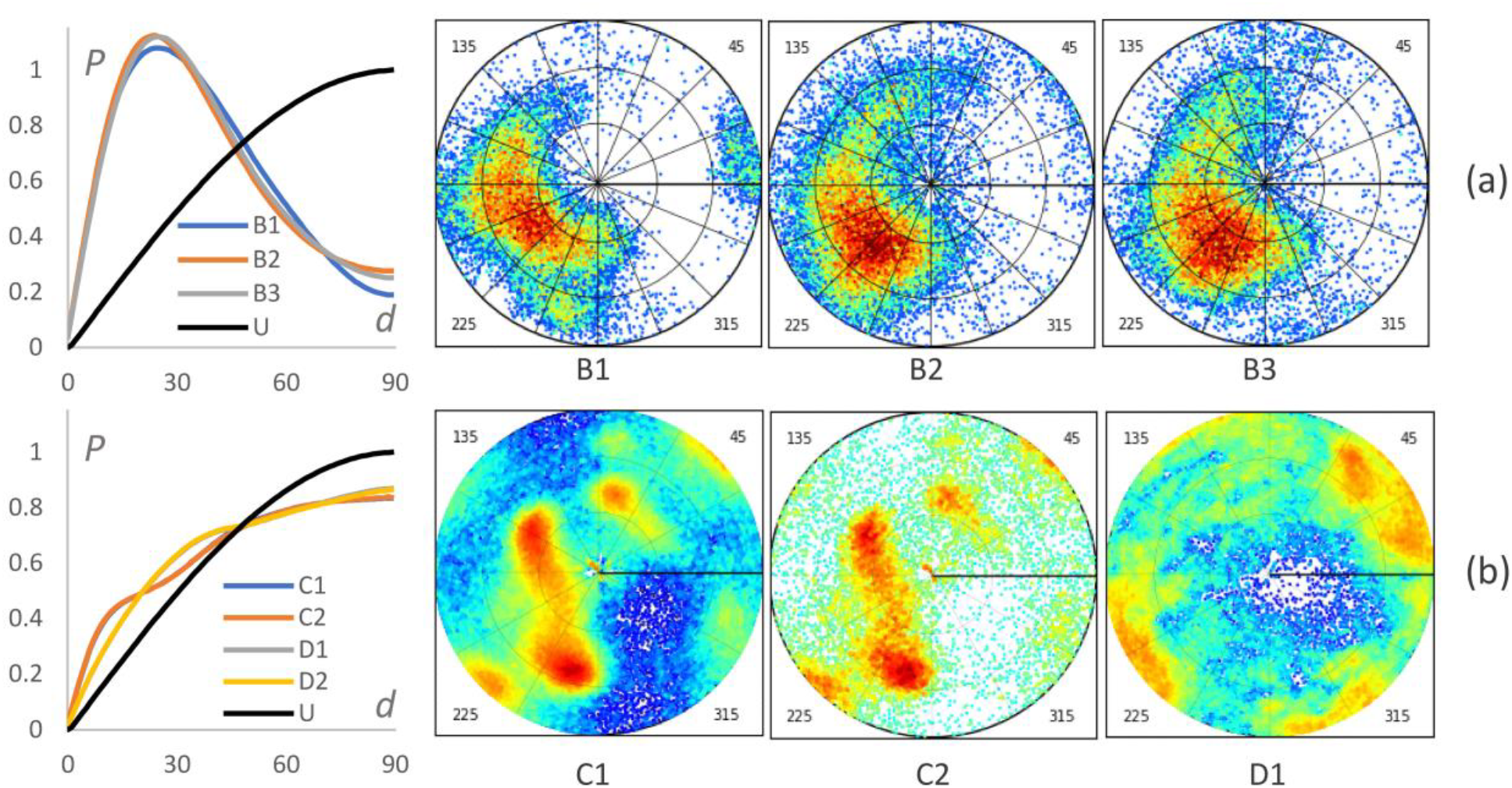
Frequency curves and 2D views diagrams for several subsets of the data sets B, C and D (Table 1 and main text for their formal definitions). Black curve (U) is theoretical for the uniform views distribution.

Next, we tested the robustness of the results against reducing of the number of views. For this purpose, we used experimental data from two additional structures, labeled C and D. Their respective projection sets, C1 and D1, contained a large number of views, approximately 140,000 for C1 and about 74,000 for D1. We then selected smaller subsets, C2 and D2, composed roughly 10% and 20% of the original projections, respectively (**Table 2**) and analyzed all four data sets.

The frequency curves for the subsets were practically indistinguishable from those of the full sets (**Fig. 4b**), consistent with the analysis of the simulated distributions. The differences between the statistical characteristics of the full sets and their subsets were on the order of 10^−3^. The curve for C1 (and C2) exhibites two peaks, one more pronounced and one less, indicating the presence of two clusters of views. In contrast, the D1 (and D2) curves display a single broader peak. Notably, the C1 curve is much closer to the theoretical uniform distribution curve than those observed for structure B; this is even more pronounced for D1. The statistical measures Δ*P*, σ*P*, and ΔC, support these observations (**Table 2**).

Indeed, the 2D views diagrams (**Fig. 4**) corroborate the data quality assessments inferred from the frequency curves.

### 3.2. Severely non-uniformly distributed sets

Next, we examined how the frequency curves and numerical characteristics reflect improvements in cryo-EM projection data sets that were initially severely non-uniformly distributed.

Set E contained about 600,000 projections. For analysis, we randomly selected a subset, E1, of approximately 29,000 projections (**Table 2**). The calculated frequency curve clearly reveals the presence of overrepresented views (**Fig. 5a**, left diagram). Using the program *VUE*, we automatically reduced these overrepresented views, first yielding set E2 with about 28,000 projections, then further to set E3 with about 17,000 projections. The corresponding curves show the progressive approach toward a uniform distribution. Finally, using the available *VUE* option (Urzhumtseva *et al*., 2024; Barchet *et al*., prepared for submission), we completed the references to the underrepresented views of set E3, producing set E4 (about 28,000 projections). The resulting curve is indistinguishable from the theoretical uniform-distribution curve, in agreement with the statistical characteristics (**Table 1**).

**Figure 5.**
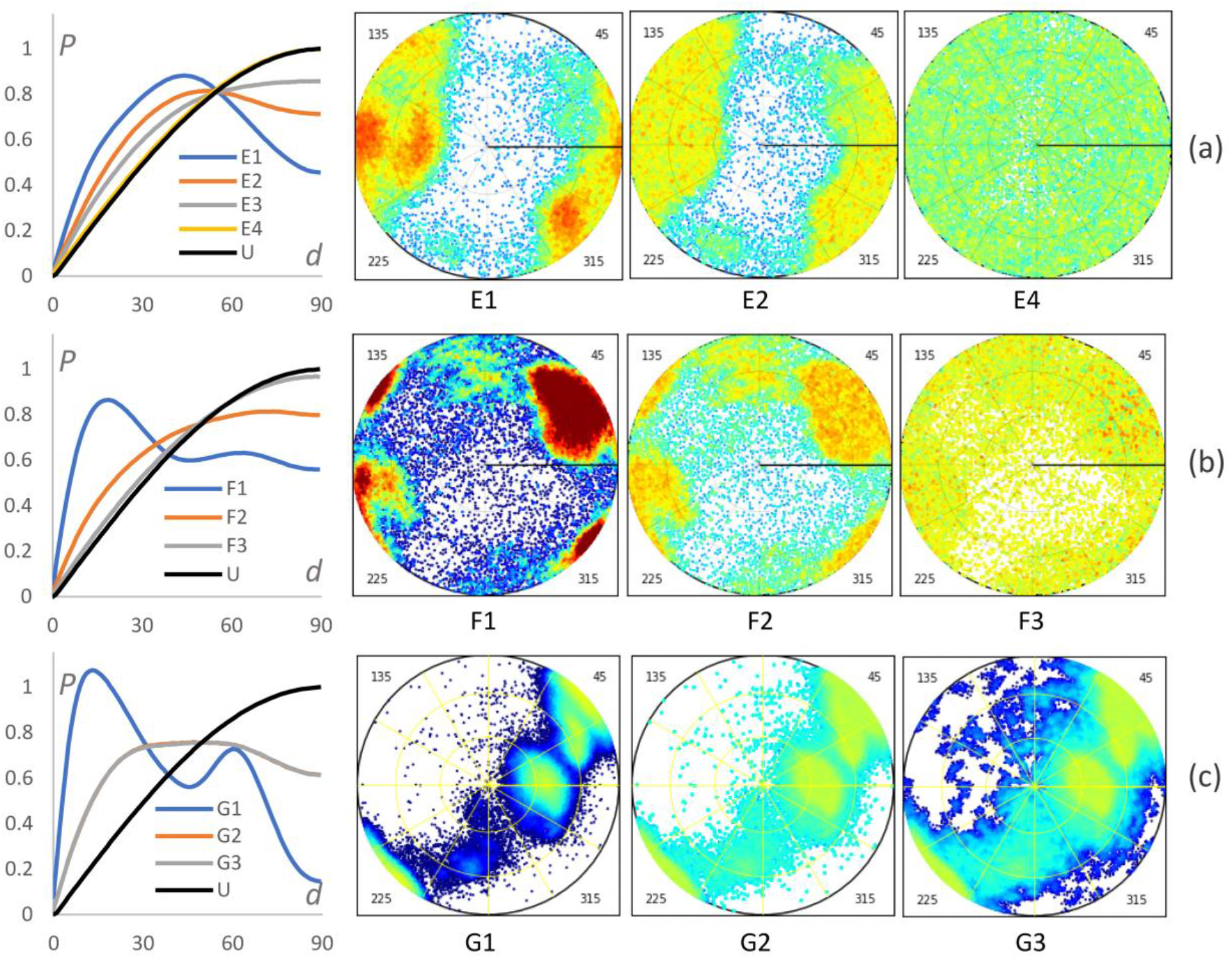
Frequency curves and 2D views diagrams for several subsets of the data sets E, F and G (Table 1 and main text for their formal definitions). Black curve (U) is theoretical for the uniform views distribution.

The corresponding 2D views projections fully agree with the curve analysis, except that the initial data set E1 actually contained three clusters positioned close to each other, rather than a single cluster (**Fig. 5a**).

Data set F1 contained about 45,000 projections, which we analyzed in full (**Table 2**). The frequency curve reveals a dominant cluster of views, accompanied by a smaller cluster located about 60–70 degrees away. Using the program *VUE*, we proportionally reduced the overrepresented views (set F2), resulting in a distribution that was already close to uniform according to the distance frequencies. The curve for data set F3, obtained after completing the most underrepresented views of F2, is practically indistinguishable from the theoretical curve for a uniform distribution. The corresponding 2D views diagrams **(Fig 5b**) fully confirm these observations.

The data set G contained more than 100,000 projections. For our analysis, we consistently worked with representative subsets containing about 10% of the full data sets. The curve for the G1 subset (∼ 11,500 projections) clearly reveals two sharp clusters and no projections at a large angular distance from them, *i.e*., indicating empty regions (**Fig. 5c**, left diagram). To balance the views distribution while keeping approximately the same total number of projections, we proportionally removed overrepresented views and supplemented underrepresented ones, following a procedure similar to that used to obtain E4 and F3. In generating such sets, we also applied a perturbation option that randomly modified the orientations of artificially generated copies of the underrepresented views. This was intended to improve Fourier space coverage, even though the Fourier coefficients derived from such perturbed projections were necessarily inexact (analysis of such procedure is beyond the scope of this work and is given in Barchet *et al*., prepared for submission; its results are not important for the current work). Completing the underrepresented views when applying a mean angular perturbation of one degree (set G2) significantly improved the distribution, as indicated by both the frequency curve and the statistical parameters. However, it still fell short of a truly uniform distribution. Increasing the mean shift to an impractically large value of 20 degrees (set G3) did not yield further improvement, which can be explained if the underrepresented views were adjacent to empty regions. Indeed, the 2D views distributions for these sets confirm this hypothesis (**Fig. 5c**).

### 3.3. Particular case

Data set H1 contained more than 337,000 projections. For analysis, we randomly selected a subset, H2, consisting of about 5% of the views. Unlike all previous cases, the frequency curve for this data set exhibites a ‘jumping’ behavior (**Fig. 6**). The curve calculated for the entire set H1 was identical, indicating that the unusual pattern is not due to subset selection. We also ruled out the possibility that this behavior resulted from merging two separate runs, sets H3 and H5. While the curves for H3 and H5 are differed from each other, each exhibites the same ‘jumping’ characteristic (**Fig. 6**). Such behavior can be explained if the views are distributed on a kind of grid (Appendix A).

**Figure 6.**
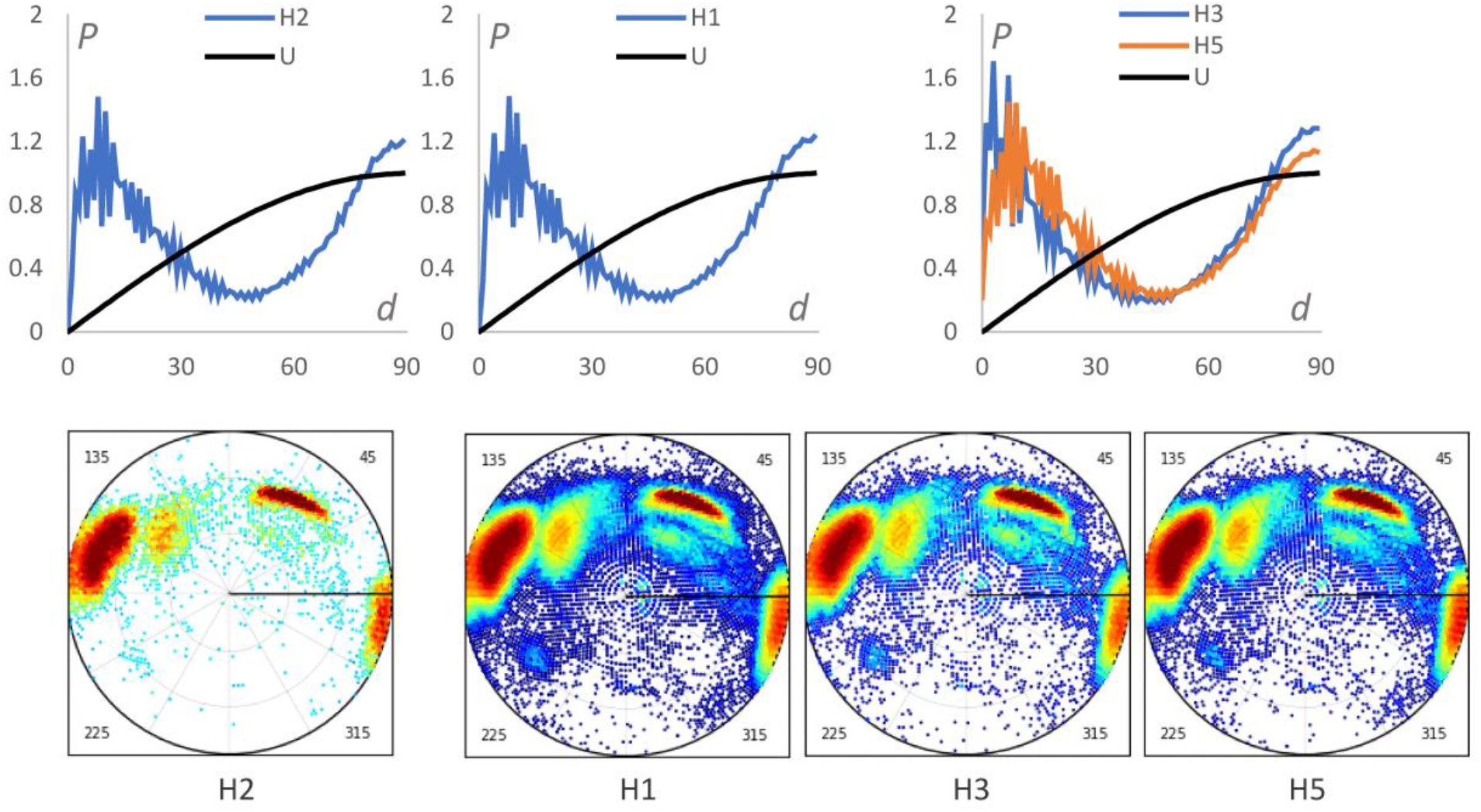
Frequency curves and 2D views diagrams for several subsets of the data set H (Table1 and main text for their formal definitions). Black curve (U) is theoretical for the uniform views distribution.

Indeed, the 2D diagrams reveal that the points representing views are arranged in rows and circles. This data set H1 corresponds to an initial projection matching step without subsequent refinement of the projection orientations. Interestingly, the frequency curves reveal differences between subsets H3 and H5 that are barely noticeable in the 2D diagrams.

### 3.4. Analysis of the measures

Among the proposed numerical characteristics, some provide overlapping information or are less practical for interpretation. **Fig. 7** presents the cumulative curves C(*d*), for the experimental data sets and their modified versions, as we calculated them previously for the simulated data **(Fig. 3**). Compared to the frequency curves P(*d*). the cumulative curves appear less convenient for discerning details of the views distribution. Their main advantage is inherent smoothness by definition. However, this same property can conceal important dataset features, as illustrated by set H. Another benefit is their near insensitivity to the number of bins. We concluded that the maximal point-by-point difference, ΔC between the calculated and theoretical cumulative functions may be more informative than other integrative characteristics, such as σ*P*, while the curves C(*d*) themselves may be less required than P(*d*).

**Figure 7.**
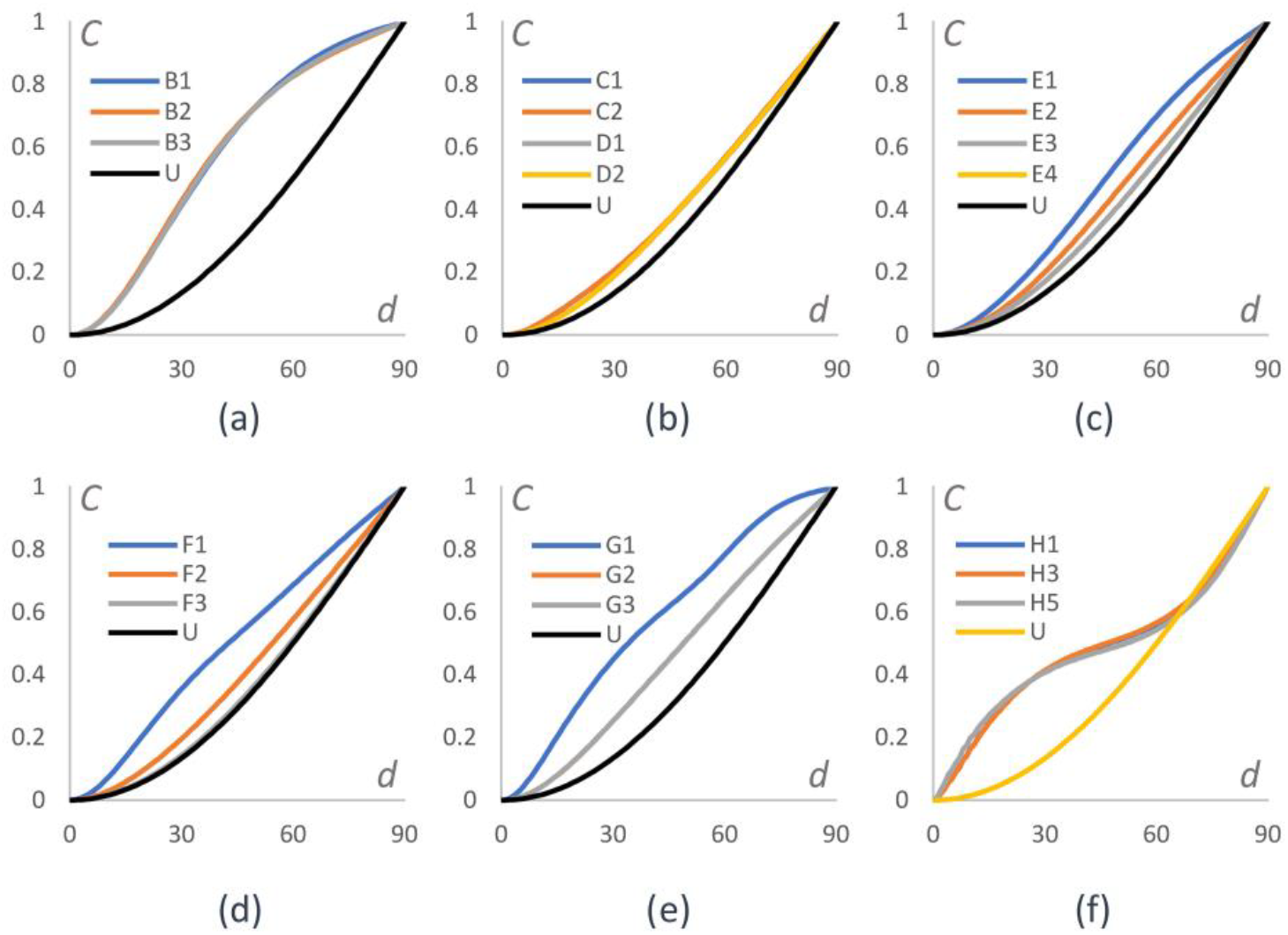
Cumulative functions for the data sets B - H. Black curve (U) stands for the theoretical function for the uniformly distributed set.

Comparison of the characteristics σP and ΔC for the experimental data sets analyzed above shows a high correlation, with an almost linear dependence. A noticeable deviation from this trend was observed only for the particular case of data set H (**Fig. 8**). The same **Fig. 8** also demonstrates that the point-by-point measure Δ*P* and the integrative measure σ*P* are similarly well correlated, with a significant departure from near-linearity again occurring only for data set H.

**Figure 8.**
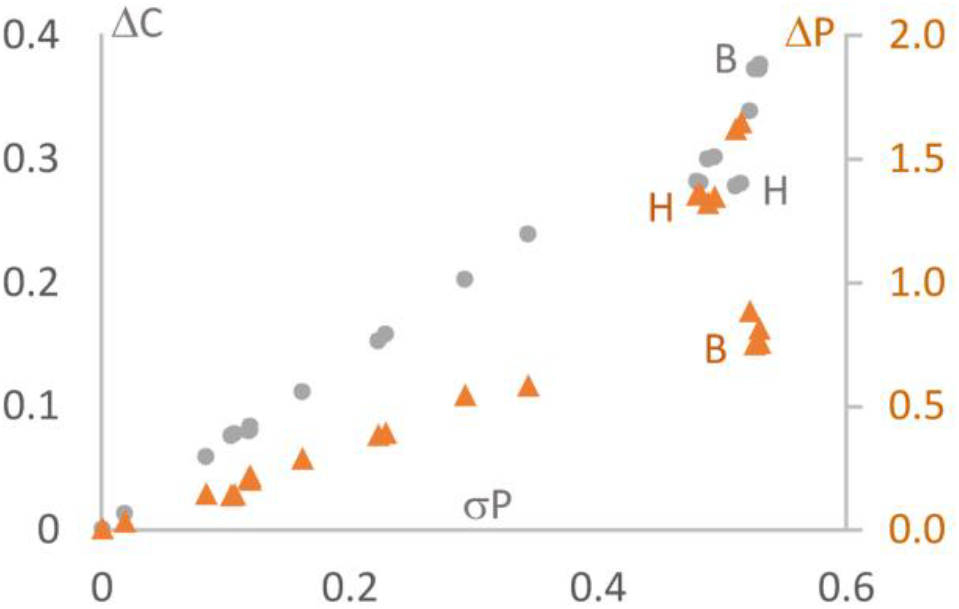
Common distribution of σP, ΔP and ΔC for experimental data sets and their modifications.

In summary, the combination of the frequency curve P(*d*) together with the values of ΔC and Δ*P* provides a robust quantitative description of how much a views distribution deviates from the uniform case. In all situations examined in this work, these metrics effectively replace the need for 2D views distribution diagrams.

## 4. Computations and program

The frequency curves for analyzing orientations of 2D projections, as defined in this work, together with the corresponding numerical characteristics, can be computed using the *NUViews* program (Non-Uniformly distributed Views analysis). is written in standard Python 3 and relies only on the NumPy and MatPlotLib libraries. It is a stand-alone tool that requires no installation. The input files describing projection orientations are compatible with a variety of common formats and software packages in the field, for example, the star format (*Relion*, Scheres, 2012), hed format (*Imagic*, van Heel *et al.*, 1996) or files created by *cryoSPARC* (Punjani *et al*., 2017). The analysis does not require access to the projection images themselves.

The program calculates the histogram of geodesic angular distances between points on the spherical surface corresponding to the view directions. This histogram is then converted into a probability distribution and plotted alongside the theoretical curve for a uniform distribution for direct comparison. The resulting values are also saved in a text file, enabling users to plot and compare curves for multiple data sets using external software, such as Excel, PowerPoint or LibreOffice. The measures of ΔP and ΔC are computed and reported both in the log file and on the plot showing the P_*calc*_(θ) and *P*_*theor*_(θ) curves. **Fig. 9b** gives an example.

**Figure 9.**
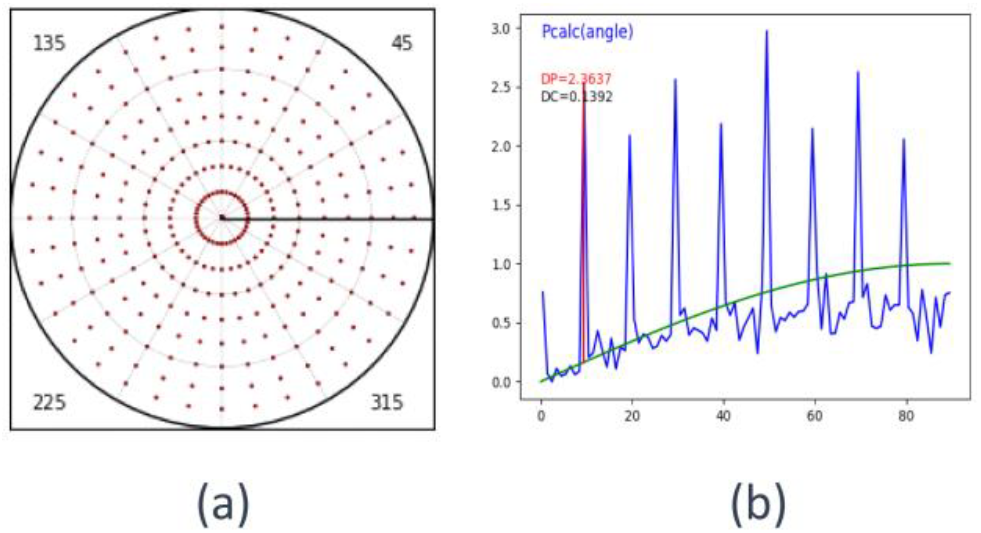
Simulated data set of the projections generated on a regular angular grid. a) 2D-diagram prepared by the program *VUE*; Urzhumtseva *et al*., 2024); b) respective frequency curve compared with the theoretical curve for the uniform distribution (diagram generated by the program *NUViews* described in this work).

The computational cost scales with the square of the number of projections. To save time, the program allows random, proportional selection of a subset of any specified size from the input projections. If desired, this subset can be saved in the same format as the original input file.

While the calculations could be greatly accelerated by rewriting the program in C++ or Fortran, we chose to retain Python for its flexibility and ease of use, and instead optimized the core histogram calculation algorithm. The most CPU-intensive step is computing the dot product of two vectors and then taking its arccosine to obtain the required distance. Since the arccosine function is monotonic, we first compute the histogram of the dot product values directly and convert it to a distance histogram only at the final step. This approach yields nearly an order-of-magnitude reduction in computing time. To minimize conversion errors, the dot product histogram is computed using a fine grid: by default, 90,000 bins are used when the corresponding angular histogram contains 90 bins, though this value, affecting the CPU time rather marginally, can be adjusted through the input parameters.

The program, along with an example, is freely available upon request from the author and from the site https://git.cbi.igbmc.fr/sacha/nuviews-quantify-views-distribution.

## 5. Discussion

Traditionally, the distribution of 2D projections in cryo-EM is analyzed using spatial or 2D diagrams. This visual representation has clear advantages, as color-based displays can be highly informative. However, projecting a spherical surface onto a plane is non-trivial, and much of the information can be inaccessible to color-blind users. Such diagrams are also less convenient for archiving in databases, and they do not easily allow numerical comparison between different structures.

To address these limitations, we propose compressing the 2D distribution information into frequency curves, and further reducing these curves to one or a few numerical values. Calculations with both simulated and experimental data sets illustrate that these data well characterize the views distribution. Each single plot in **Figs. 2, 4, 5, 6, 9** illustrates well main features of a few respective 2D-diagrams. Moreover, in case of the data set H, such curve allowed to identify two groups of projections merged in a single set that the 2D-diagrams did not show.

The proposed approach is not intended to replace the 2D projection diagrams, but rather to complement them by simpler, quantitative descriptors. In particular, the measures ΔC (8) and Δ*P* (10) can be incorporated into ‘Table 1’ commonly reported in structural publications and into cryo-EM databases such as the EMDB to provide a concise description of the angular distribution of particle views. The data bases can also include the plot of *P*_*calc*_(θ) similar to other plots currently implemented.

## Acknowledgments

The author thanks C. Barchet, L. Fréchin, R. Bahena Ceron, Y.-C. Chen and O. von Loeffelholz, for sharing their experimental data, and B.P. Klaholz for important discussion and his interest to the project.

This work, especially in its experimental part, was supported by CNRS, Agence Nationale pour la Recherche (ANR), the Interdisciplinary Thematic Institute IMCBio, as part of the ITI 2021-2028 program of the University of Strasbourg, CNRS and Inserm, was supported by IdEx Unistra (ANR-10-IDEX-0002) and by SFRI-STRAT’US project (ANR 20-SFRI-0012), EUR IMCBio (ANR-17-EURE-0023) under the framework of the France 2030 program and LabexNetRNA (ANR-10-LABX-0036_NETRNA) administered by ANR. The electron microscope facility was supported by the Region Grand Est, FEDER, the French Infrastructure for Integrated Structural Biology (FRISBI) ANR-10-INBS-0005 / France 2030 program, EquipEx^+^ France-Cryo-EM (ANR-21-ESRE-0046) and Instruct-ERIC.

## Appendix A

To highlight the effect observed for the experimental set H, we prepared a file describing a set of views corresponding to a coarse angular grid on a sphere. **Fig. 9** shows the respective 2D diagram and the frequency curve.

